# Evaluation of population structure inferred by principal component analysis or the admixture model

**DOI:** 10.1101/2023.06.06.543934

**Authors:** Jan van Waaij, Song Li, Genís Garcia-Erill, Anders Albrechtsen, Carsten Wiuf

## Abstract

Principal component analysis (PCA) is commonly used in genetics to infer and visualize population structure and admixture between populations. PCA is often interpreted in a way similar to inferred admixture proportions, where it is assumed that individuals belong to one of several possible populations or are admixed between these populations. We propose a new method to assess the statistical fit of PCA (interpreted as a model spanned by the top principal components) and to show that violations of the PCA assumptions affect the fit. Our method uses the chosen top principal components to predict the genotypes. By assessing the covariance (and the correlation) of the residuals (the differences between observed and predicted genotypes), we are able to detect violation of the model assumptions. Based on simulations and genome wide human data we show that our assessment of fit can be used to guide the interpretation of the data and to pinpoint individuals that are not well represented by the chosen principal components. Our method works equally on other similar models, such as the admixture model, where the mean of the data is represented by linear matrix decomposition.

## Introduction

Principal component analysis (PCA) and model-based clustering methods are popular ways to disentangle the ancestral genetic history of individuals and populations. One particular model, the admixture model (Pritchard *et al*. 2000), has played a prominent role because of its simple structure and, in some cases, easy interpretability. PCA is often seen as being model free but as noted by Engelhardt and Stephens (2010), the two approaches are very similar. The interpretation of the results of a PCA analysis is often based on assumptions similar to those of the admixture model, such that admixed individuals are linear combinations of the eigenvectors representing unadmixed individuals. In this way, the admixed individuals lie in-between the unadmixed individuals in a PCA plot. As shown for the admixture model, there are many demographic histories that can lead to the same result (Lawson *et al*. 2018a) and many demographic histories that violate the assumptions of the admixture model (Garcia-Erill and Albrechtsen 2020). As we will show, this is also the case for PCA, since it has a similar underlying model (Engelhardt and Stephens 2010). In fact just as in the admixture model, the choice of the number of PCs is similar to choosing the number of ancestral components in the admixture model and, if this choice is too low, the PCA will not reflect the correlation between individuals truthfully.

The admixture model states that the genetic material from each individual is composed of contributions from *k* distinct ancestral homogeneous populations. However, this is often contested in real data analysis, where the ancestral population structure might be much more complicated than that specified by the admixture model. For example, the *k* ancestral populations might be heterogeneous themselves, the exact number of ancestral populations might be difficult to assess due to many smaller contributing populations, or the genetic composition of an individual might be the result of continuous migration or recent backcrossing, which also violates the assumptions of the admixture model. Furthermore, the admixture model assumes individuals are unrelated, which naturally might not be the case. This paper is concerned with assessing the fit of PCA building on the special relationship with the admixture model (Engelhardt and Stephens 2010). In particular, we are interested in quantifying the model fit and assessing the validity of the model at the level of the sample as well as at the level of the individual. Using real and simulated data we show that the fit from a PCA analysis is affected by violations of the admixture model.

We consider genotype data *G* from *n* individuals and *m* SNPs, such that *G*_*si*_ ∈ {0, 1, 2} is the number of reference alleles for individual *i* and SNP *s*. Typically, *G*_*si*_ is assumed to be binomially distributed with parameter Π_*si*_, where Π_*si*_ depends on the number of ancestral populations, *k*, their admixture proportions and the ancestral population allele frequencies. For clustering based analysis such as ADMIXTURE (Alexander and Lange 2011), *k* is the number of clusters used to explain the data, while in PCA, the *k −*1 top principal components are used to explain the data. We give the specifics of the admixture model in the next section and show its relationship to PCA in ‘Materials and methods’.

Several methods aim to estimate the best *k* in some sense (Alexander and Lange 2011; Evanno *et al*. 2005; Pritchard *et al*. 2000; Raj *et al*. 2014; Wang 2019), but finding such *k* does not imply the data fit the model (Lawson *et al*. 2018b; Janes *et al*. 2017). In statistics, it is standard to use residuals and distributional summaries of the residuals to assess model fit (Box *et al*. 2005). The residual of an observation is defined as the difference between the observed and the predicted value (estimated under some model). Visual trends in the residuals (for example, differences between populations) are indicative of model misfit, and large absolute values of the residuals are indicative of outliers (for example due to experimental errors, or kinship). If the model is correct, a histogram of the residuals is expected to be mono-modal centered around zero (Box *et al*. 2005).

In our context, Garcia-Erill and Albrechtsen (2020) argue that trends in the residual correlation matrix carries information about the underlying model and might be used for visual model evaluation. A method is designed to assess whether the correlation structure agrees with the proposed model, in particular, whether it agrees with the proposed number of homogeneous ancestral populations (Garcia-Erill and Albrechtsen 2020). However, even in the case the model is correctly specified, the residuals are in general correlated (Box *et al*. 2005), and therefore, trends might be observed even if the model is true, leading to incorrect model assessment. To adjust for this correlation, a leave-one-out procedure, based on maximum likelihood estimation of the admixture model parameters, is developed that removes the correlation between residuals in the case the model is correct, but not if the model is mis-specified (Garcia-Erill and Albrechtsen 2020). This approach could also be applied to PCA, where expected genotypes could be calculated using probabilistic PCA (Meisner *et al*. 2021). This leave-one-out procedure is, however, computationally expensive.

To remedy the computational difficulties, we take a different approach to investigate the correlation structure. We suggest two different ways of estimating the true correlation matrix of the residuals. The first is simply the empirical correlation matrix of the residuals. The second is an estimate of the true correlation matrix under a given model (that is, given the number of ancestral populations). Both are simple to compute. Under mild regularity assumptions, these two estimators agree if the model is correct and the number of SNPs is large. Hence, their difference is expected to be close to zero, when the admixture model is not violated. If the difference is considerably different from zero, this is proof of model misfit.

To explore the adequacy of the proposed method, we investigate different ways to calculate the predicted values of the genotype (hence, the residuals), using Principal Component Analysis (PCA) in different ways. However, we also show that this approach can be used on estimated admixture proportions. Specifically, we use 1) an uncommon but very useful PCA approach (here, named PCA 1) based on unnormalized genotypes (Cabreros and Storey 2019; Chen and Storey 2015), 2) PCA applied to mean centred data (PCA 2), see Patterson *et al*. (2006), and 3) PCA applied to mean and variance normalised data (PCA 3) (Patterson *et al*. 2006). All three approaches are computationally fast and do not require separate estimation of ancestral allele frequencies and population proportions, as in Garcia-Erill and Albrechtsen (2020). Calculating the residuals from PCA or ADMIXTURE output involves a single matrix multiplication, which for most data sets only takes seconds (or minutes for very large data sets) and in all cases is much faster than performing the PCA or ADMIXTURE analysis itself. Hence, the calculation of the residuals is computationally inexpensive. Additionally, we show that this approach can also be applied to output from, for example, the software ADMIXTURE (Alexander *et al*. 2009) to estimate Π_*si*_ for each *s* and *i*, and to calculate the residuals from these estimates. An overview of PCA can be found in Jolliffe and Cadima (2016).

We demonstrate that our proposed method works well on simulated and real data, when the predicted values (and the residuals) are calculated in any of the four mentioned ways. Furthermore, we back this up mathematically by showing that the two correlation measures agree (if the number of SNPs is large) under the correct admixture model for PCA 1 and PCA 2. For the latter, a few additional assumptions are required. The estimated covariance matrix (and correlation matrix) under the proposed model might be seen as a correction term for population structure. Subtracting it from the empirical covariance matrix, thus gives a covariance estimate with baseline zero under the correct model, independent of the population structure. It is natural to suspect that similar can be done in models with population structure and kinship, which we will pursue in a subsequent study.

In the next section, we describe the model, the statistical approach to compute the residuals, and how we evaluate model fit. In addition, we give mathematical statements that show how the method performs theoretically. In ‘Results’, we provide analysis of simulated and real data, respectively. We end with a discussion.

## Materials and methods

### Non-technical overview

We provide a non-technical overview that perhaps allows the reader to skip the technical parts of ‘Materials and methods’ and jump directly to ‘Results’. The residuals are commonly studied quantities in statistics to assess goodness-of-fit of a model. It is defined as the difference between the observed and the predicted value. In our context, this would be 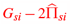, where *G*_*si*_ is the observed genotype of SNP *s* in individual *i*, and 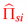 its predicted value, assuming *G*_*si*_ is a binomial random variable with parameter Π_*si*_, and 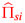 an estimator of Π_*si*_. Rather than studying the residuals themselves, we study the correlation of residuals between individuals, which captures, in case of model misfit, trends between populations and between individuals (for example due to kinship). The predicted values might be calculated in one of several ways. We advocate estimates obtained by PCA and suggest three different versions (PCA 1, PCA 2, PCA 3), where PCA 2 is standard PCA on mean normalized data, PCA 3 is standard PCA on mean and variance normalized data, and PCA 1 is based on Chen and Storey (2015). The discrepancies between the methods are not important and can be glossed over, as the three methods seem to perform in similar ways. To assess model fit, we compute an estimate of the true correlation of residuals in two different way (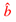 and *ĉ*), and demonstrate that these agree if the model is correct (provide a good fit), and otherwise not.

### Notation

For an *ℓ*_1_ ×*ℓ*_2_ matrix *A* = (*A*_*i j*_)_*i, j*_, *A*_*⋆i*_ denotes the *i*-th column of *A, A*_*i⋆*_ the *i*-th row, *A*^*T*^ the transpose matrix, and rank(*A*) the rank. The Frobenius norm of a square *ℓ* × *ℓ* matrix *A* is

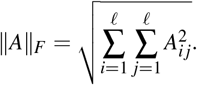

A square matrix *A* is an orthogonal projection if *A*^2^ = *A* and *A*^*T*^ = *A*. A symmetric matrix has *n* real eigenvalues (with multiplicity) and the eigenvectors can be chosen such that they are orthogonal to each other. If the matrix is positive (semi-)definite, then the eigenvalues are positive (non-negative).

For a random variable/vector/matrix *X*, its expectation is denoted 𝔼 [*X*] (provided it exist). The variance of a random variable *X* is denoted var(*X*), and covariance between two random variables *X, Y* is denoted cov(*X, Y*) (provided they exist). Similarly, for a random vector *X* = (*X*_1_, …, *X*_*n*_), the covariance matrix is denoted cov(*X*). For a sequence *X*_*m*_, *m* = 0, …, of random variables/vectors/matrices, if *X*_*m*_ → *X*_0_ as *m* → ∞ almost surely (convergence for all realisations but a set of zero probability), we leave out ‘almost surely’ and write *X*_*m*_ → *X*_0_ as *m* → ∞ for convenience.

### The PCA and the admixture model

We consider a model with genotype observations from *n* individuals, and *m* biallelic sites (SNPs), where *m* is assumed to be (much) larger than *n, m*≥ *n*. The genotype *G*_*si*_ of SNP *s* in individual *i* is assumed to be a binomial random variable

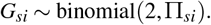

In matrix notation, we have *G*∼ binomial(2, Π) with expectation 𝔼 (*G* |Π) = 2Π, where *G* and Π are *m*× *n* dimensional matrices. Conditional on Π, we assume the entries of *G* are independent random variables.

Furthermore, we assume the matrix Π takes the form Π = *FQ*, where *Q* is a (possibly unconstrained) *k* ×*n* matrix of rank *k* ≤*n*, and *F* is a (possibly unconstrained) *m*× *k* matrix, also of rank *k* (implying Π likewise is of rank *k*, Lemma 13). Entry-wise, this amounts to

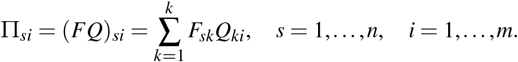

For the binomial assumption to make sense, we must require the entries of Π to be between zero and one.

In the literature, this model is typically encountered in the form of an admixture model with *k* ancestral populations, see for example, Pritchard *et al*. (2000); Garcia-Erill and Albrechtsen (2020). The general unconstrained setting which applies to PCA has also been discussed (Cabreros and Storey 2019). In the case of an admixture model, *Q* is a matrix of ancestral admixture proportions, such that the proportion of individual *i*’s genome originating from population *j* is *Q*_*ji*_. Furthermore, *F* is a matrix of ancestral SNP frequencies, such that the frequency of the reference allele of SNP *s* in population *j* is *F*_*s j*_. In many applications, the columns of *Q* sum to one.

While we lean towards an interpretation in terms of ancestral population proportions and SNP frequencies, our approach does not enforce or assume the columns of *Q* (the admixture proportions) to sum to one, but allow these to be unconstrained. This is advantageous for at least two reasons. First, a proposed model might only contain the major ancestral populations, leaving out older or lesser defined populations. Hence, the sum of ancestral proportions might be smaller than one. Secondly, when fitting a model with fewer ancestral populations than the true model, one should only require the admixture proportions to sum to at most one.

### The residuals

Our goal is to design a strategy to assess the hypothesis that Π is a product of two matrices. As we do not know the true *k*, we suggest a number *k*′ of ancestral populations and estimate the model parameters under this constraint. That is, we assume a model of the form

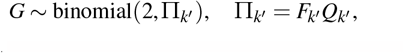

where each entry of *G* follows a binomial distribution. *Q*_*k*_′ has dimension *k*′ × *n, F*_*k*_′ has dimension *m* × *k*′, and rank(*Q*_*k*_′) = rank(*F*_*k*_′) = *k*′, hence also rank(Π_*k*_′) = *k*′. Throughout, we use the index *k*′ to indicate the imposed rank condition, and assume *k*′ ≤*k* unless otherwise stated.

The latter assumption is only to guarantee the mathematical validity of certain statements, and is not required for practical use of the method. Our approach is build on the residuals, the difference between observed and predicted data. To define the residuals, we let *P* : ℝ^*n*^→ ℝ^*n*^ be the orthogonal projection onto the *k*-dimensional subspace spanned by the *k* rows of (the true) *Q*, hence *P* = *Q*^*T*^ (*QQ*^*T*^)^−1^*Q*, and *QP* = *Q* (note that *QQ*^*T*^ is invertible by assumption on the rank of *Q*). Let 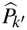 be an estimate of *P* based on the data *G*, and assume 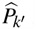 is an orthogonal projection onto a *k*′-dimensional subspace. Later in this section, we show how an estimate 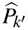 can be obtained from an estimate of *Q*_*k*_′ or an estimate of Π_*k*_′. Estimates of these parameters might be obtained using existing methods, based on, for example, maximum likelihood analysis (Wang 2003; Alexander *et al*. 2009; Garcia-Erill and Albrechtsen 2020). Furthermore, for the three PCA approaches, an estimate of the projection matrix can simply be obtained from eigenvectors of a singular value decomposition (SVD) of the data matrix.

We define the *m* × *n* matrix of residuals by

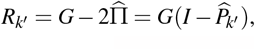

where *G* is the observed data and 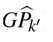 the predicted values. The latter might also be considered an estimate of 2Π, the expected value of *G*. This definition of residuals is in line with how the residuals are defined in a multilinear regression model as the difference between the observed data (here, *G*) and the projection of the data onto the subspace spanned by the regressors (here, 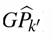). The essential difference being that in a multilinear regression model, the regressors are known and does not depend on the observed data, while 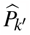 is estimated from the data.

We assess the model fit by estimating the true covariance (correlation) matrix of the residuals in two ways. We will show that these two estimators of the true covariance (correlation) matrix are approximately identical, if the number of SNPs is large and the model is correct, while they are expected to differ if the model is not correct. By comparing the two estimators, we thus have a measure of model fit. To begin with, we quantify the true covariance matrix of the residuals as the covariance matrix of *T*_*s⋆*_ = *G*_*s⋆*_ (*I*− *P*) (which by the modelling assumptions is independent of the SNP *s*), that is, the covariance of *R*_*s⋆*_ with the estimated projection matrix replaced by the true projection matrix,

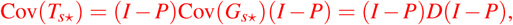

where we have used (3), *QP* = *Q*, and *D* is the diagonal matrix with entries

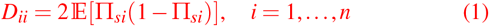

(the genotype variance of individual *i* as *G*_*si*_ is binomial).

The first estimator we consider is the *empirical covariance matrix* 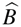 with entries

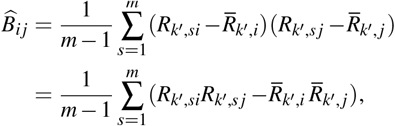

where

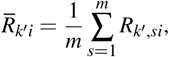

and the corresponding *empirical correlation matrix* with entries

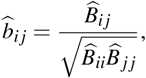

*i, j* = 1, …, *n*. Secondly, we consider the *estimated covariance matrix* under the proposed model,

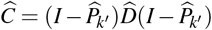

with corresponding *estimated correlation matrix*,

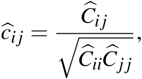

*i, j* = 1, …, *n*. Here, 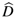 is the *n* ×*n* diagonal matrix containing the average heterozygosities of each individual,

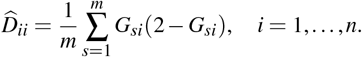

It follows from Lemma 4 in the appendix, that 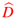 converges to *D* as *m*→ ∞. Thus, 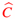 might be considered an estimate of the true covariance matrix.

Under reasonable regularity conditions, we can quantify the behaviour of 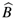 and 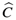 as the number of SNPs become large. Specifically, we assume the rows of *F* are independent and identically distributed with distribution Dist(*μ*, ∑), where *μ* denote the *k*-dimensional mean vector of the distribution, and ∑ the *k* × *k*-covariance matrix, that is,

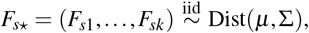

*s* = 1, …, *m*. The matrix *Q* is assumed to be non-random, that is, fixed. These assumptions are standard and typically used in simulation of genetic data, see for example, Pickrell and Pritchard (2012); Cabreros and Storey (2019); Garcia-Erill and Albrechtsen (2020). Often dist(*μ*, ∑) is taken to be the product of *k* independent uniform distributions in which case *μ* = 0.5(1, 1, …, 1) and ∑ is a diagonal matrix with entries 1/12, though other choices have been applied, see for example Balding and Nichols (1995); Conomos *et al*. (2016).

The proofs of the statements are in the appendix.

#### Theorem 1.

*Let k*′ ≤ *k. Under the given assumptions, suppose further that* 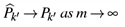, *for some matrix P*_*k*_′. *Then, P*_*k*_′ *is an orthogonal projection. Furthermore, the following holds*,

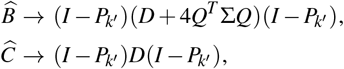

*as m* → ∞. *Hence, also*

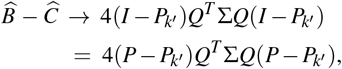

*as m* →∞. *For k*′ = *k, if P*_*k*_ = *P, then the right hand side is the zero matrix, whereas this is not the case in general for k*′ *< k*.

#### Theorem 2.

*Assume k*′ = *k and P*_*k*_ = *P. Furthermore, suppose as in Theorem 1 and that the vector with all entries equal to one is in the space spanned by the rows of Q (this is, for example, the case if the admixture proportions sum to one for each individual). Then*,

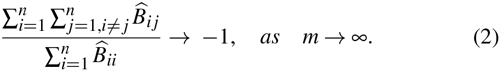

*In addition, if Q takes the form*

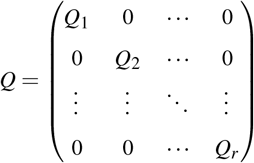

*where Q*_*ℓ*_ *has dimension* 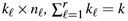 *and* 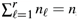, *then* (2) *holds for each component of n*_*ℓ*_ *individuals. If Q*_*ℓ*_ = (1 … 1), *then*

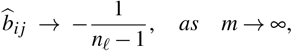

*for all individuals i, j in the ℓ-th component, irrespective the form of Q*_*ℓ*_′, *ℓ*′ ≠ *ℓ*.

#### Theorem 3.

*Assume k*′ = *k and P*_*k*_ = *P. Furthermore, suppose as in Theorem 1 and that Q takes the form*

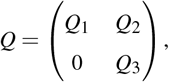

*where Q*_1_ = (1 … 1) *has dimension* 1 × *n*_1_, *n*_1_ ≤ *n. Then*, 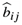 *converges as m* → ∞ *to a value larger than or equal to* 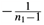, *for all i, j* = 1, …, *n*_1_.

The same statements in the last two theorems hold with 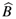 and 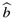 replaced by 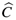 and 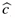, respectively.

The three theorems provide means to evaluate the model. In particular, Theorem 1 might be used to assess the correctness (or appropriateness) of the proposed *k*′, while Theorem 2 and Theorem 3 might be used to assess whether data from a group of individuals (e.g., a modern day population) originates from a single ancestral population, irrespective of the origin of the remaining individuals. We give examples in ‘Results’.

The work flow is shown in Algorithm 1. We process real and simulated genotype data using PCA 1, PCA 2, PCA 3, and the software ADMIXTURE, and evaluate the fit of the model.

#### Algorithm 1

Work flow of the proposed method

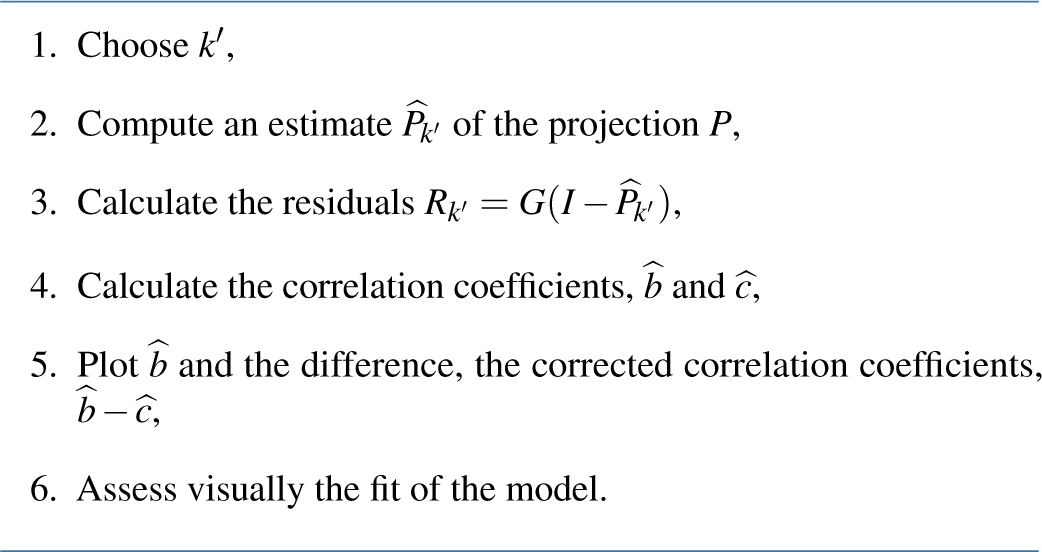

### Estimation of *P*_*k*_′

Estimation of *Q, F*, and Π has received considerable interest in the literature, using for example, maximum likelihood (Wang 2003; Alexander *et al*. 2009), Bayesian approaches (Pritchard *et al*. 2000) or PCA (Engelhardt and Stephens 2010).

We discuss different ways to obtain an estimate 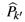 of *P*.

#### ***Using an estimate*** 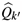 ***of*** *Q*_*k*_′

An estimate 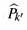 might be obtained by projecting onto the subspace spanned by the *k*′ rows of 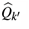,

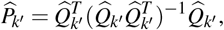

assuming rank 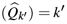 for the calculation to be valid.

We apply this approach to estimate the projection matrix using output from the software ADMIXTURE.

#### ***Using an estimate*** 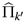 ***of*** Π_*k*_′

Let 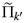 be *k*′ linearly independent rows chosen from 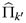 (out of *m* rows). Then, an estimate 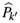 of *P*_*k*_′ is

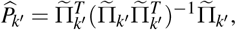

assuming rank 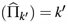 for the calculation to be valid. Alternatively, one might apply the Gram-Schmidt method in which case the vectors are orthonormal by construction and 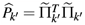. The estimate 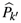 is independent of the choice of the *k*′ rows, provided rank 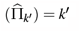.

#### Using PCA 1

We consider a PCA approach, originally due to Chen and Storey (2015), to estimate the space spanned by the rows of *Q*. We follow the procedure laid out in Cabreros and Storey (2019).

Let 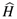 be the symmetric matrix

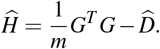

Since 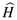 is symmetric, all eigenvalues are real and the matrix is diagonalisable. Furthermore, 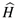 is a variance adjusted version of 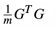, see (1). Let *u*_1_, …, *u*_*k*_′ be *k*′≤ *k* orthogonal eigenvectors belonging to the *k*′ largest eigenvalues of 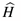, counted with multiplicities. Define the *n* ×*k*′ matrix *U*_*k*_′ = (*u*_1_, …, *u*_*k*_′) and the *n* ×*n* orthogonal projection matrix

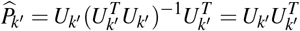

onto the subspace given by the span of the vectors *u*_1_, …, *u*_*k*_′.

In Appendix B we show convergence and validity results for PCA 1. For *k*′ = *k*, the correct row space of *Q* is found eventually, but not *Q* itself. If *k*′ *< k*, then a subspace of this row space is found, corresponding to the *k*′ largest eigenvalues. As the data is not mean centred, we discard the first principal component, and use the subsequent *k*′ −1 eigenvectors and eigenvalues. The principal components are named PC0, PC1, …, such that comparison to PCA 2 and PCA 3 is straightforward (by discarding PC0).

#### Using PCA 2 (mean centred data)

A popular approach to estimation of Π in the admixture model is PCA based on mean centred data, or mean and variance normalised data (Pritchard *et al*. 2000; Engelhardt and Stephens 2010; Patterson *et al*. 2006).

Let 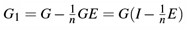 be the SNP-wise mean centred genotypes, where *E* is an *n*× *n* matrix with all entries equal to one. Following the exposition and notation in Cabreros and Storey (2019), let *G*_1_ = *U* Δ*V*^*T*^ be the SVD of *G*_1_, where Δ*V*^*T*^ consists of the rowwise principal components of *G*_1_, ordered according to the singular values. Define

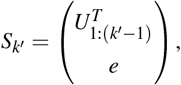

where *e* = (11 … 1) is a vector with all entries one, and 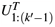 contains the top *k*′ − 1 rows of *U*^*T*^. Then, an estimate of the projection

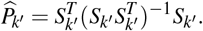

Under certain additional assumptions, PCA 2 comes with the same mathematical guarantees as PCA 1 about convergence and validity, see Appendix B for details.

#### Using PCA 3 (mean and variance normalised data)

Let *G*_2_ = *W* ^−1^*G*_1_ be the SNP mean and variance normalised genotypes, where *W* is an *m*′ ×*m*′ diagonal matrix with *s*-th entry being the observed standard deviation of the genotypes of SNP *s*. All SNPs for which no variation are observed are removed, hence the number of SNPs might be smaller than the original number, *m*′≤ *m*. Following the same procedure as for PCA 2, let *G*_2_ = *U* Δ*V*^*T*^ be the SVD of *G*_2_, where Δ*V*^*T*^ consists of the row-wise principal components of *G*_2_, ordered according to the singular values. Define

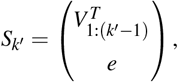

where *e* = (11 … 1), and 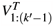 contains the top *k*′ − 1 rows of *V*^*T*^. Then, an estimate of the projection is 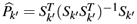.

We are not aware of any theoretical justification of this procedure similar to Theorem 1, but it appears to perform well in many practical situations, according to our simulations.

### Simulation of genotype data

We simulated genotype data from different demographic scenarios using different sampling strategies. We deliberately choose different sampling strategies to challenge the method. We first made simple simulations that illustrate the problem of model fit as well as to demonstrate the theoretical and practical properties of the residual correlations that arise from having data from a finite number of individuals and a large number of SNPs. An overview of the simulations are given in Table 1.

**Table 1.** Overview of simulations.

| Scenario | $k$ | $n$ | $m$ | Description | $F_{is}^a$ |
| --- | --- | --- | --- | --- | --- |
| 1 | 3 | 20,20,20 | 500K | Unadmixed | Unif(0,1) |
| 1 | 3 | 10,20,30 | 500K | Unadmixed | Unif(0,1) |
| 2 | 2 | 20,20,20 | 500K | Admixed | Unif(0,1) |
| 2 | 2 | 10,20,30 | 500K | Admixed | Unif(0,1) |
| 3 | | 500 | 100K<sup>b</sup> | Spatial with $F_{st} = 0.001$<br>between adjacent populations | Unif(0.01,0.99) |
| 4 | 4 | 50,50,50,50,0<sup>c</sup> | 10M<sup>d</sup> | Ghost admixture | Beta(0.3,0.3) |
| 5 | 2 | 20,20,50 | 500K | Recent hybrids | Unif(0.05,0.95) |

<sup>a</sup> Ancestral allele frequencies, $i = 1, \dots, k$ <sup>b</sup> after applying MAF > 5% filtering, 88,082 remained.<sup>c</sup> No individuals are sampled from the ghost population.<sup>d</sup> after applying MAF > 5% filtering, 694,285 remained.

In the first two scenarios, the ancestral allele frequencies are simulated independently for each ancestral population from a uniform distribution, *F*_*si*_ ∼Unif(0, 1) for each site *s* = 1, …, *m* and each ancestral population *i* = 1, …, *k*. In scenario 1, we simulated unadmixed individuals from three populations with either an equal or an unequal number of sampled individuals from each population. In scenario 2, we simulated two ancestral populations and a population that is admixed with half of its ancestry coming from each of the two ancestral populations.

In scenario 3, we set *F*_*si*_ ∼Unif(0.01, 0.99) and simulated spatial admixture in a way that resembles a spatial decline of continuous gene flow between populations living in a long narrow island. We first simulated a single population in the middle of the long island. From both sides of the island, we then recursively simulated new populations from a Balding-Nichols distribution (Balding and Nichols 1995) with parameter *F*_*st*_ = 0.001 using the R package ‘bnpsd’ (Ochoa and Storey 2019). In this way, each pair of adjacent populations along the island has an *F*_*st*_ of 0.001. Additional details on the simulation and an schematic visualization can be found in Figure 2 of Garcia-Erill and Albrechtsen (2020).

In scenario 4, we first simulated allele frequencies for an ancestral population from a symmetric beta distribution with shape parameter 0.03, *F*_*si*_ ∼Beta(0.3, 0.3), which results in an allele frequency spectrum enriched for rare variants, mimicking the human allele frequency spectrum. We then sampled allele frequencies from a bifurcating tree (((pop1:0.1,popGhost:0.2):0.05,pop2:0.3):0.1,pop3:0.5), where pop1 and popGhost are sister populations and pop3 is an outgroup. Using the Balding-Nichols distribution and the *F*_*st*_ branch lengths of the tree (see Figure 5), we sampled allele frequencies in the four leaf nodes. Then, we created an admixed population with 30% ancestry from popGhost and 70% from pop2. We sampled 10 million genotypes for 50 individuals from each population except for the ghost population which was not included in the analysis, and subsequently removed sites with a sample minor allele frequency below 0.05, resulting in a total of 694,285 sites.

In scenario 5, we simulated an ancestral population with allele frequencies from a uniform distribution *F*_*si*_∼ Unif (0.05, 0.95), from which we sampled allele frequencies for two daughter populations from a Balding-Nichols distribution with *F*_*st*_ = 0.3 from the ancestral population, using ‘bnpsd’. We then created recent hybrids based on a pedigree where all but one founder has ancestry from the first population. The number of generations in the pedigree then determines the admixture proportions and the age of the admixture where F1 individuals have one unadmixed parent from each population and backcross individuals have one unadmixed parent and the other F1. Double backcross individuals have one unadmixed parent and the other is a backcross. We continue to quadruple backcross with one unadmixed parent and the other triple backcross. Note that for the recent hybrids the ancestry of the pair of alleles at each loci is no longer independent which is a violation of the admixture model.

## Results

### Scenario 1

In this first set-up, we demonstrate the method using PCA 1 only. We simulated unadmixed individuals from *k* = 3 ancestral populations

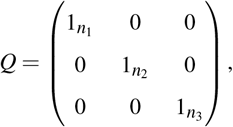

where 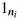 is a row vector with all elements being one, and *n*_1_ + *n*_2_ + *n*_3_ = *n*. We simulated genotypes for *n* = 60 individuals with sample sizes *n*_1_, *n*_2_ and *n*_3_, respectively, as detailed in the previous section. In Figure 1(A), we show the residual correlation coefficients for *k*′ = 2, 3 and plot the corresponding major PCs. For the PCA 1 approach, the first principal component does not relate to population structure as the data is not mean centered, and we use the following *k*′ −1 principal components. These are named PC1, PC2,…, PC(*k*′ −1).

**Figure 1.**
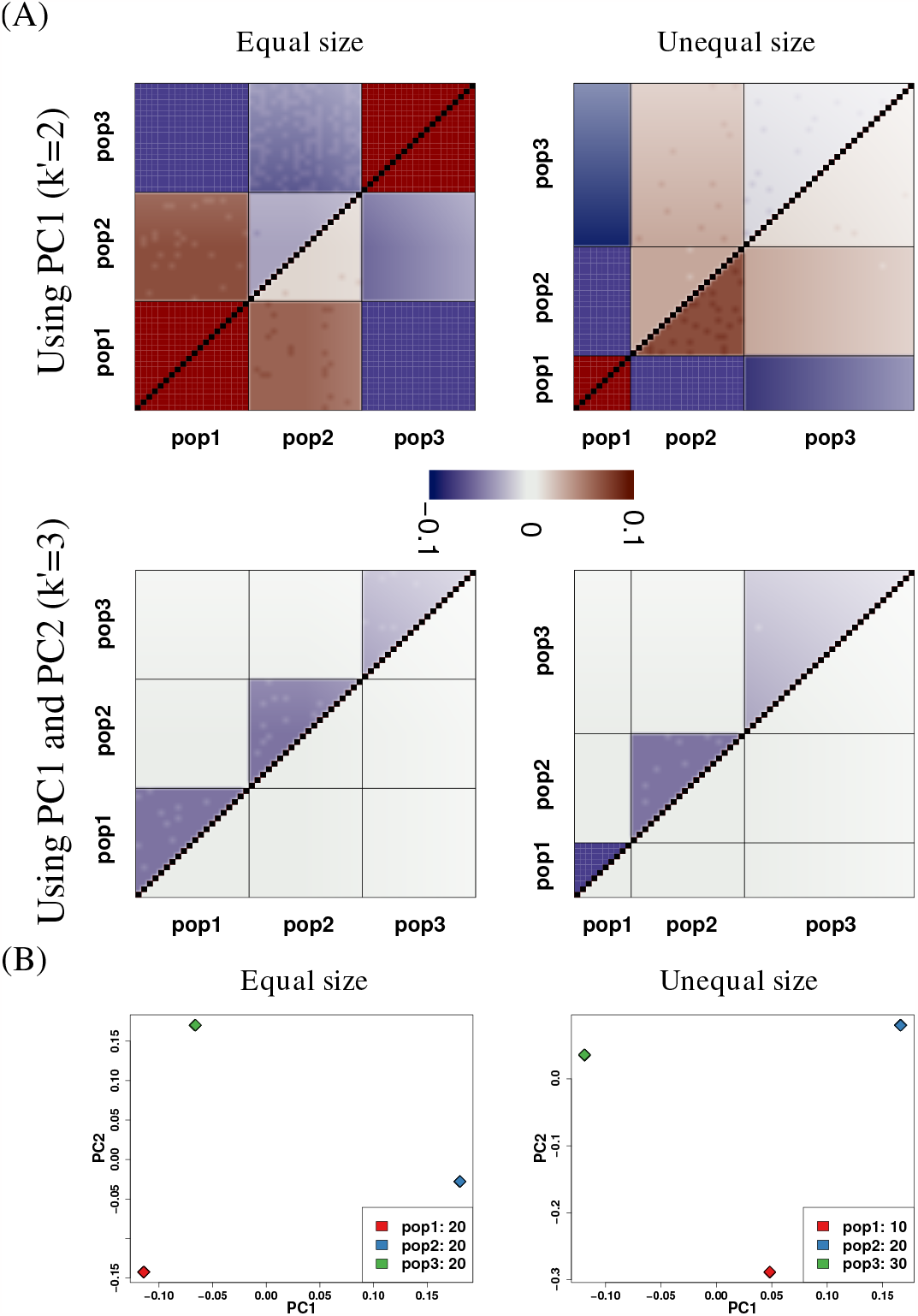
Results for simulated Scenario 1. (A) The upper triangle in the plots shows the empirical correlation coefficients 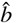 and the lower triangle shows the corrected correlation coefficients 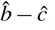. (B) The major principal components (*k*′ = 3) result in a clear separation of the three population samples (all data points within each population are almost identical).

When assuming that there are only two populations, *k*′ = 2, we note that the empirical correlation coefficients appear largely consistent within each population sample, but the corrected correlation coefficients are generally non-zero with different signs, which points to model misfit. In contrast, when assuming the correct number of populations is *k*′ = 3, the empirical correlation coefficients match nicely the theoretical values of 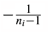, which comply with Theorem 2 (see Table 2). A fairly homogeneous pattern in the corrected correlation coefficients appears around zero across all population samples. This is a good indication that the model fits well and that the principal components PC1 and PC2 explain the data.

**Table 2.**
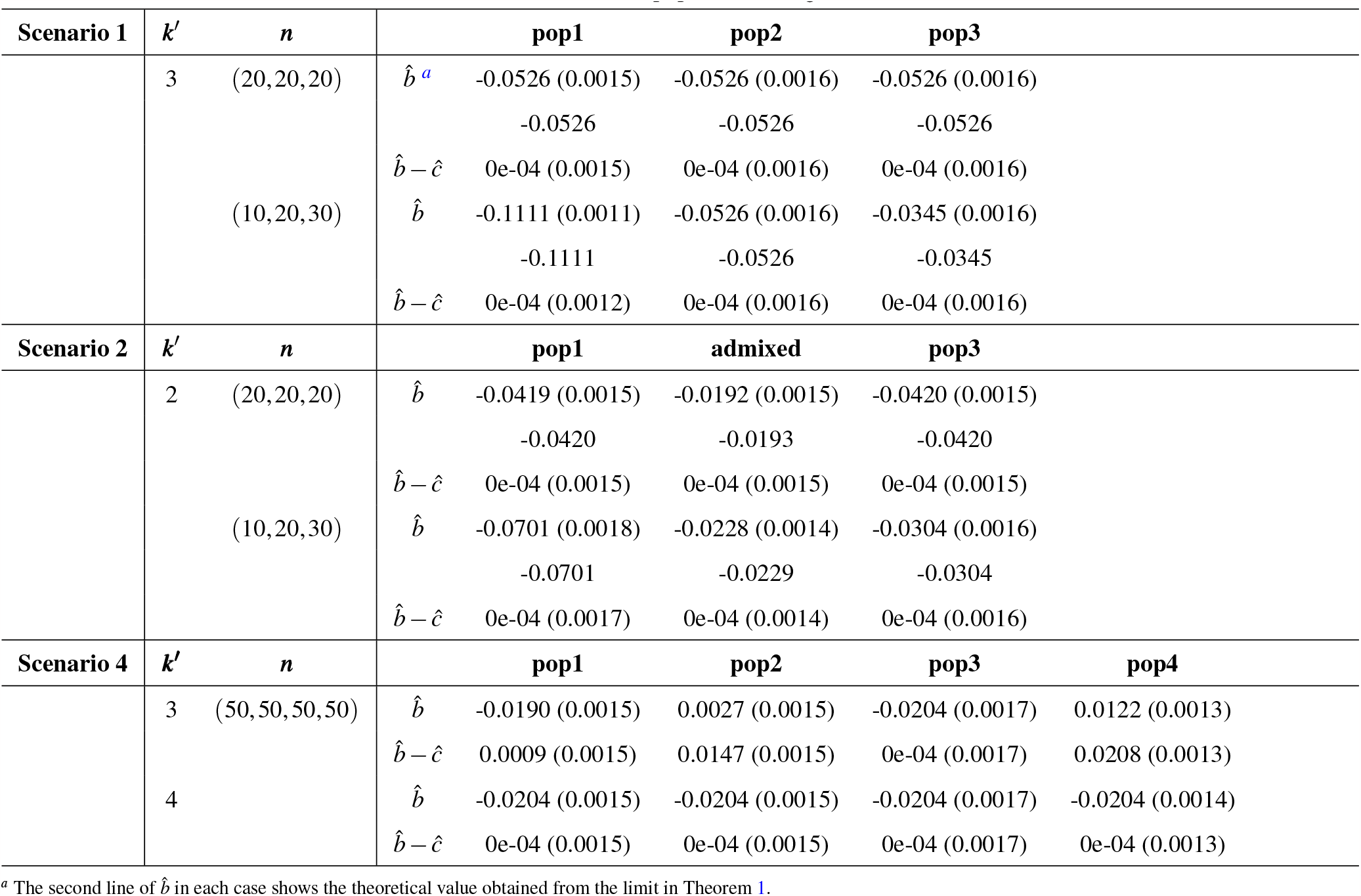
The mean (standard deviation) of 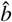 and 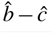 within each population using PCA 1.

### Scenario 2

In this set-up we also include admixed individuals. We simulated samples from two ancestral populations and individuals that are a mix of the two. We then applied all three PCA procedures and the software ADMIXTURE to the data. Specifically, we choose

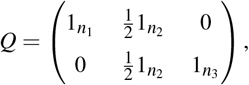

with *k* = 2 true ancestral populations, and (*n*_1_, *n*_2_, *n*_3_) = (20, 20, 20) or (*n*_1_, *n*_2_, *n*_3_) = (10, 20, 30), see the previous section for details. We analysed the data with *k*′ = 1, 2, 3, and obtained the correlation structure shown in Figures 2 and S1, and Table 2. The two standard approaches PCA 2 and PCA 3 show almost identical results, hence only PCA 2 is shown in the figures. For *k*′ = 1, none of the principal components are used and the predicted normalized genotypes is simply 0. All four methods show consistent results, in particular, for the correct *k*′ (= 2), while there are smaller discrepancies between the methods for wrong *k*′ = 1, 3. This is most pronounced for PCA 1 and ADMIXTURE. We note that the average correlation coefficient of 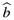 within each population sample comply with Theorem 1 (see Table 2). A fairly homogeneous pattern in the corrected correlation coefficients appears around zero across all population samples for *k*′ = 2, as in scenario 1, which shows that the model fits well. However, unlike in scenario 1, the bias for the empirical correlation coefficient is not a simple function of the sample size (see Table 2). Both *k* = 2 and *k* = 3 appear to fit the data, and higher *k* might likewise provide good fits, even though they over fit the data. In such situations, we advocate to use the smallest *k* (the most parsimoneous) that appears to provide a good fit.

**Figure 2.**
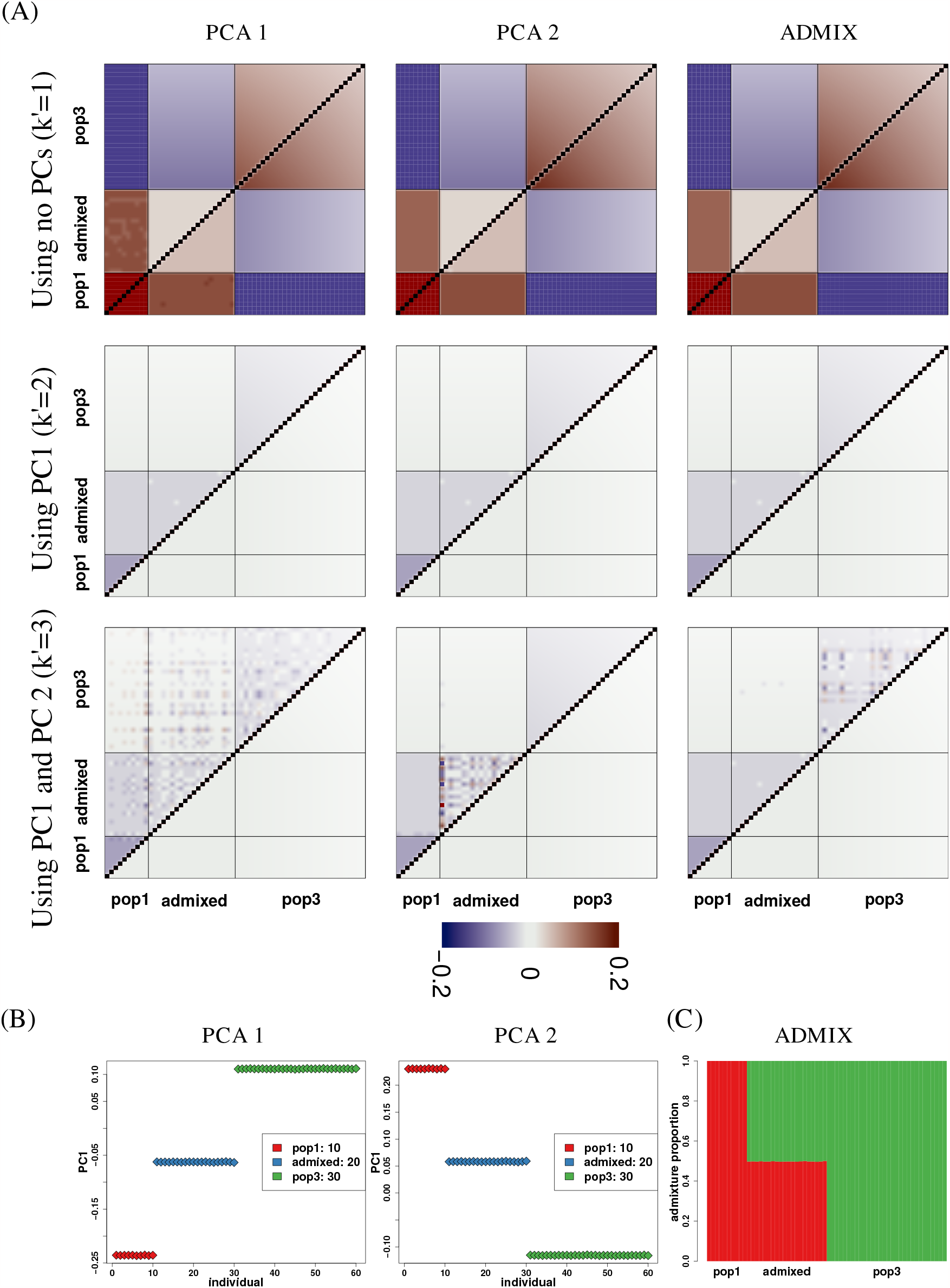
Results for simulated Scenario 2 with unequal sample sizes. (A) For each of PCA 1, PCA 2 and ADMIXTURE, the upper left triangle in the plots shows the empirical correlation 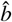 and the lower right triangle shows the difference 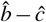 with sample sizes (*n*_1_, *n*_2_, *n*_3_) = (10, 20, 30). (B) The major principal component for the PCA based methods for *k*′ = 2 (in which case there is only one principal component). Individuals within each population sample have the same color. (C) The estimated admixture proportions in the case of ADMIXTURE.

In this case, and similarly in all other investigated cases, we don’t find any big discrepancies between the four methods. Therefore, we only show the results of PCA 1, occasionally for PCA 2, for which we have theoretical justification for the results.

### Scenario 3

We simulated genotypes for *n* = 500 individuals at *m* = 88, 082 sites with continuous genetic flow between individuals, thus there is not a true *k*. We analysed the data assuming *k*′ = 2, 3, see Figure 3. In the figure, the individuals are ordered according to the estimated proportions of the ancestral populations, hence it appears there is a color wave pattern in the empirical and the corrected correlation coefficients, see Figure 3(A). As expected, the corrected correlation coefficients are closer to zero for *k*′ = 3 than *k*′ = 2, though the deviations from zero are still large. We thus find no support for the model for either value of *k*′. This is consistent with the plots of the major PCs, that show continuous change without grouping the data into two or three clusters, see Figure 3(B). Further PCs become subsequently more noisy but still capture different patterns of continuous structure, and similarly the eigenvalues associated to each PC show a continuously declining pattern (Figure S2). See also Novembre and Stephens (2008) for a discussion of a similar phenomenon.

**Figure 3.**
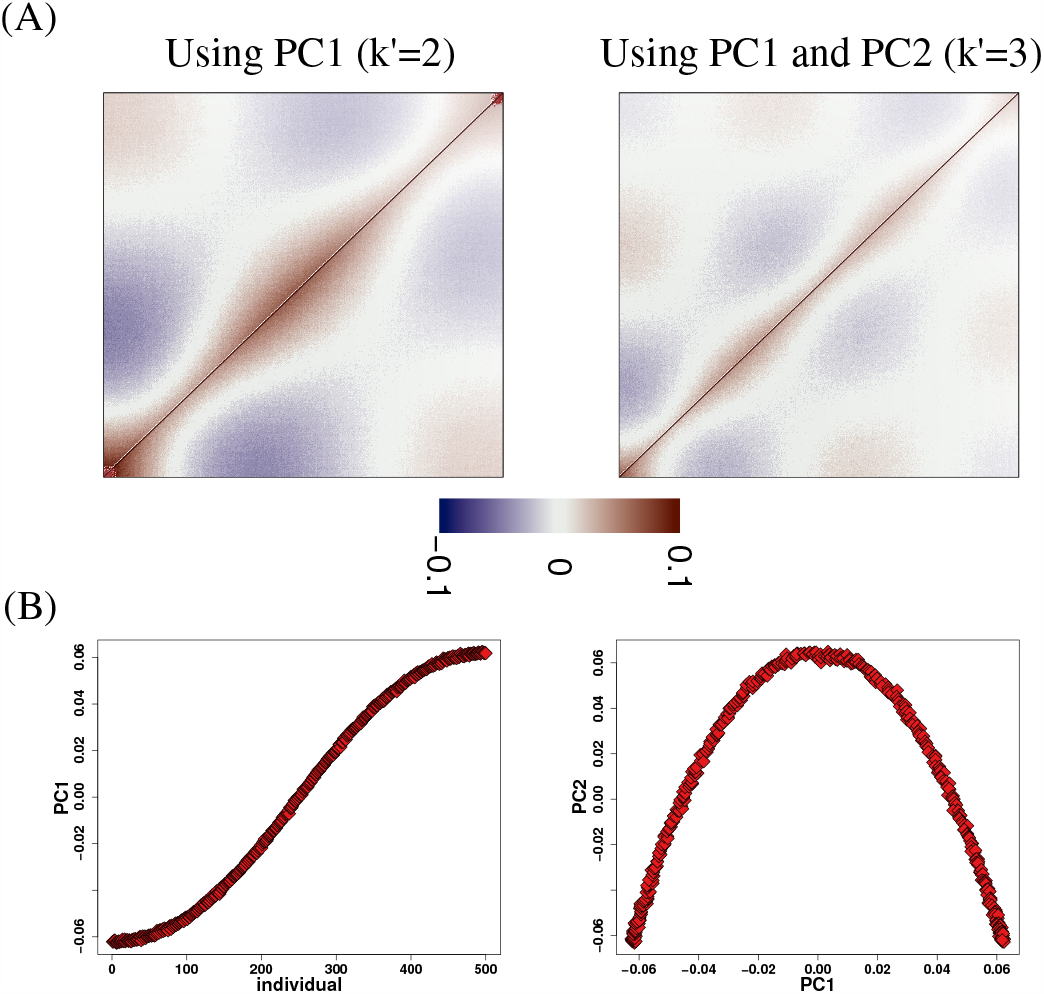
Results for simulated scenario 3. (A) The upper triangle in the plots shows the empirical correlation 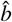 and the lower triangle shows the difference 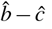. (B) The major principal components (only one in the case of *k*′ = 2).

### Scenario 4

This case is based on the tree in Figure 4(A), which includes an unsampled (so-called) ghost population, popGhost. The popGhost is sister population to pop1.

**Figure 4.**
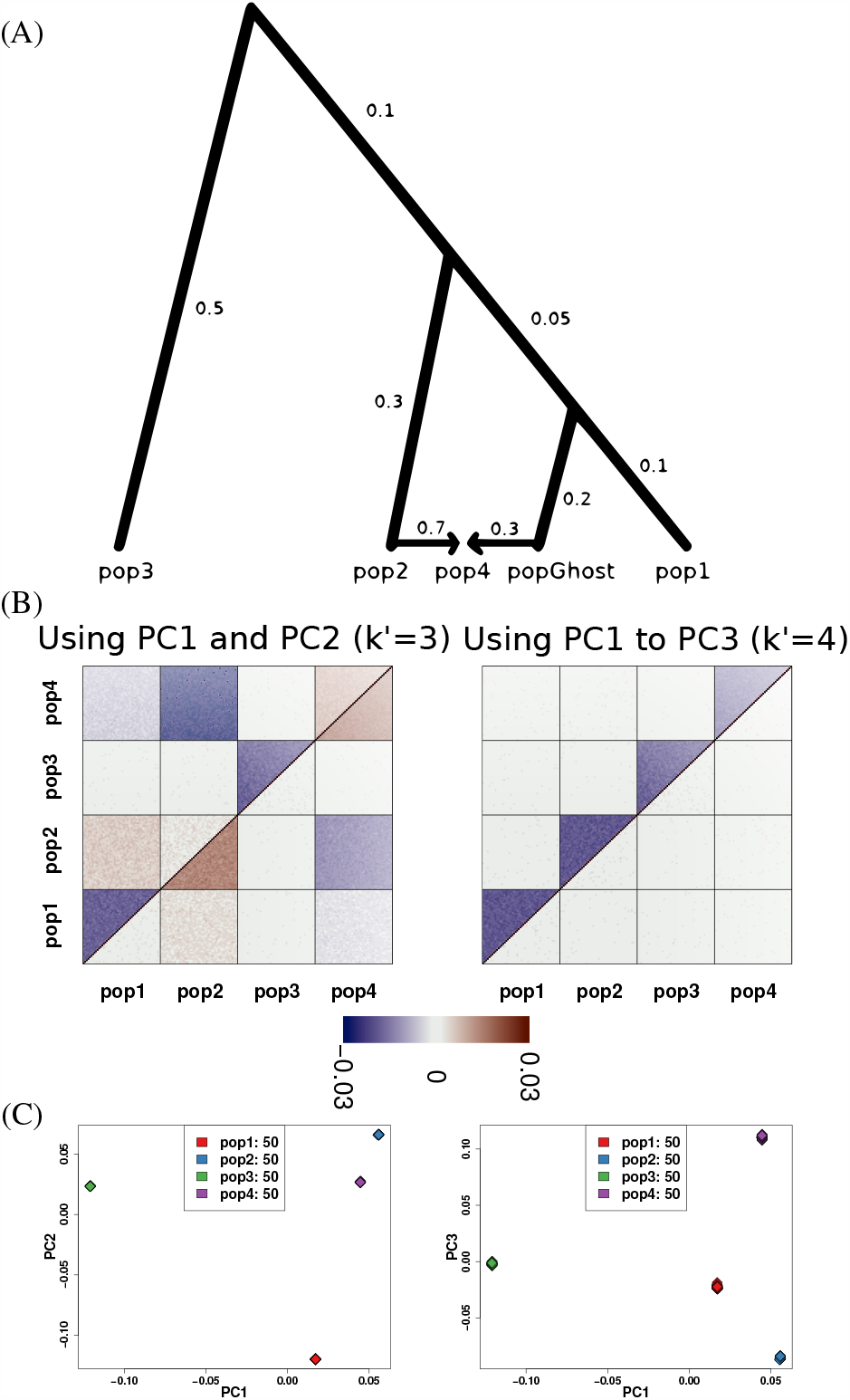
Results for simulated scenario 4. (A) Schematic of the tree used to simulate population allele frequencies, including 5 populations: pop1, pop2, pop3, pop4 and popGhost. The pop4 population is the result of admixture between pop2 and popGhost, for which there are no individuals sampled and is therefore a ghost population. The values in the branches indicate the drift in units of *F*_*ST*_. The values along the two admixture edges are the admixture proportions coming from each population. (B) The upper triangle in the plots shows the empirical correlation 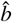 and the lower triangle shows the difference 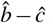. (C) The major principal components for *k*′ = 4, that result in a clear separation of the four population samples (all data points within each population sample are almost identical).

We simulated genotypes for *n* = 200 individuals: 150 unadmixed individuals from pop1, pop2, and pop3; and 50 individuals, admixed with 0.3 ancestry from popGhost and 0.7 ancestry from pop2 (as pop4), as detailed in the previous section. As there is drift between the populations and hence genetic differences, the correct *k* = 4 (pop1, pop2, pop3, popGhost). This is picked up by our method that clearly shows *k*′ = 3 is wrong with large deviation from zero in the corrected correlation coefficients. In contrast, for *k*′ = 4, the corrected correlation coefficients are almost zero (Figure 4). In this case the eigenvalues show a steep decline between PC3 and PC4, and visualizing the PCs also show that PC4 and all subsequent PCs capture only noise (Figure S3).

### Scenario 5

In the last example, we simulated two populations (originating from a common ancestral population) and created admixed populations by backcrossing, as detailed in the previous section. Thus, the model does not fulfil the assumptions of the admixture model in that the number of reference alleles are not binomially distributed, but depends on the particular backcross and the frequencies of the parental populations.

We simulate genotypes for *n* = 90 individuals at *m* = 500, 000 sites. There are 20 homogeneous individuals from each parental population, and 10 different individuals from each of the different recent admixture classes. Then, we analysed the data with *k*′ = 2 and found the corrected correlation coefficients deviated consistently from zero, in particular for one of the parental populations (Figure 5). We are thus able to say the admixture model does not provide a reasonable fit. The scree plot and visualization of subsequent PCs suggests the only PC capturing relevant information is PC1 (Figure S4). Because in this case the model violation is not due to unmodelled population substructure, approaches based on exploring the PCs approach would miss the model misfit in this scenario.

**Figure 5.**
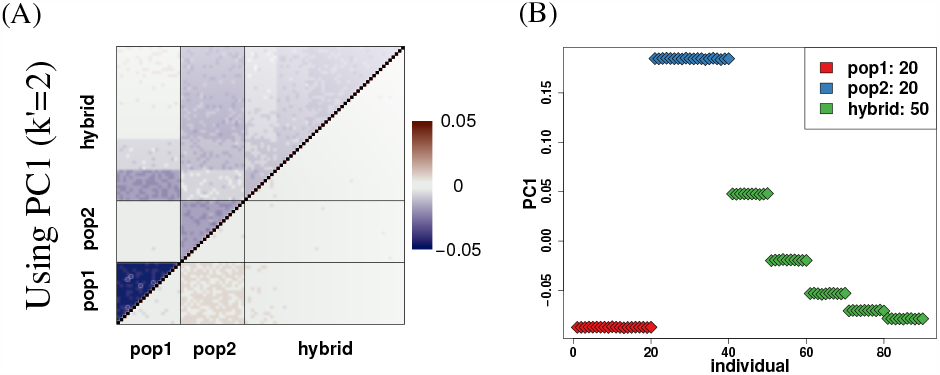
Results for simulated scenario 5 (recent admixture). (A) The upper triangle in the plots shows the empirical correlation 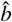 and the lower triangle shows the difference 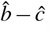. (B) The major principal component for *k*′ = 2.

### Real data

We analysed a whole genome sequencing data set from the 1000 Genomes Project (Auton *et al*. 2015), see also Garcia-Erill and Albrechtsen (2020) where the same data is used. It consists of data from five groups of different descent: a Yoruba group from Ibadan, Nigeria (YRI), residents from Southwest US with African ancestry (ASW), Utah residents with Northern and Western European ancestry (CEU), a group with Mexican ancestry from Los Angeles, California (MXL), and a group of Han Chinese from Beijing, China (CHB) with sample sizes 108, 61, 99, 64 and 103, respectively, in total, *n* = 435. We kept only sites present in the Human Origins SNP panel (Lazaridis *et al*. 2014). A total of *m* = 407, 441 SNPs were left after applying a MAF filter of 0. 05. In this example we applied PCA 2.

The results are shown in Figure 6 for *k*′ = 3, 4 with additional PC plots in Figure S5 With two PCs (*k*′ = 3) the admixed individuals, ASW and MXL, are placed between YRI, CEU and CHB (Figure 6B), suggesting that they are a mixture of three populations. However, the MXL individuals show high correlation with other MXL individuals and negative correlations with CEU and CHB individuals (Figure 6(A)), which indicates that this is not the case. The addition of PC3 (*k*′ = 4) shows that MXL have a distinct ancestry and reduces most correlation coefficients to almost zero. In Figure 6(C) each individual’s maximum correlation coefficient is shown, where high correlation indicates individuals with a bad fit. Some of these can be explained by 5 pairs of first degree relatives (Garcia-Erill and Albrechtsen 2020) and their second highest correlation coefficient is close to zero (Figure 6(D)). When plotting subsequent PCs we find that PC4 to PC8 are driven by these 5 pairs of first degree relatives from the ASW population (Figure S5). The scree plot shows a steep decline in the associated eigenvalues after PC3, after all population substructure is modelled, and a second decline after PC8 when the impact of the pairs of first degree relatives has also been accounted for (Figure S5).

**Figure 6.**
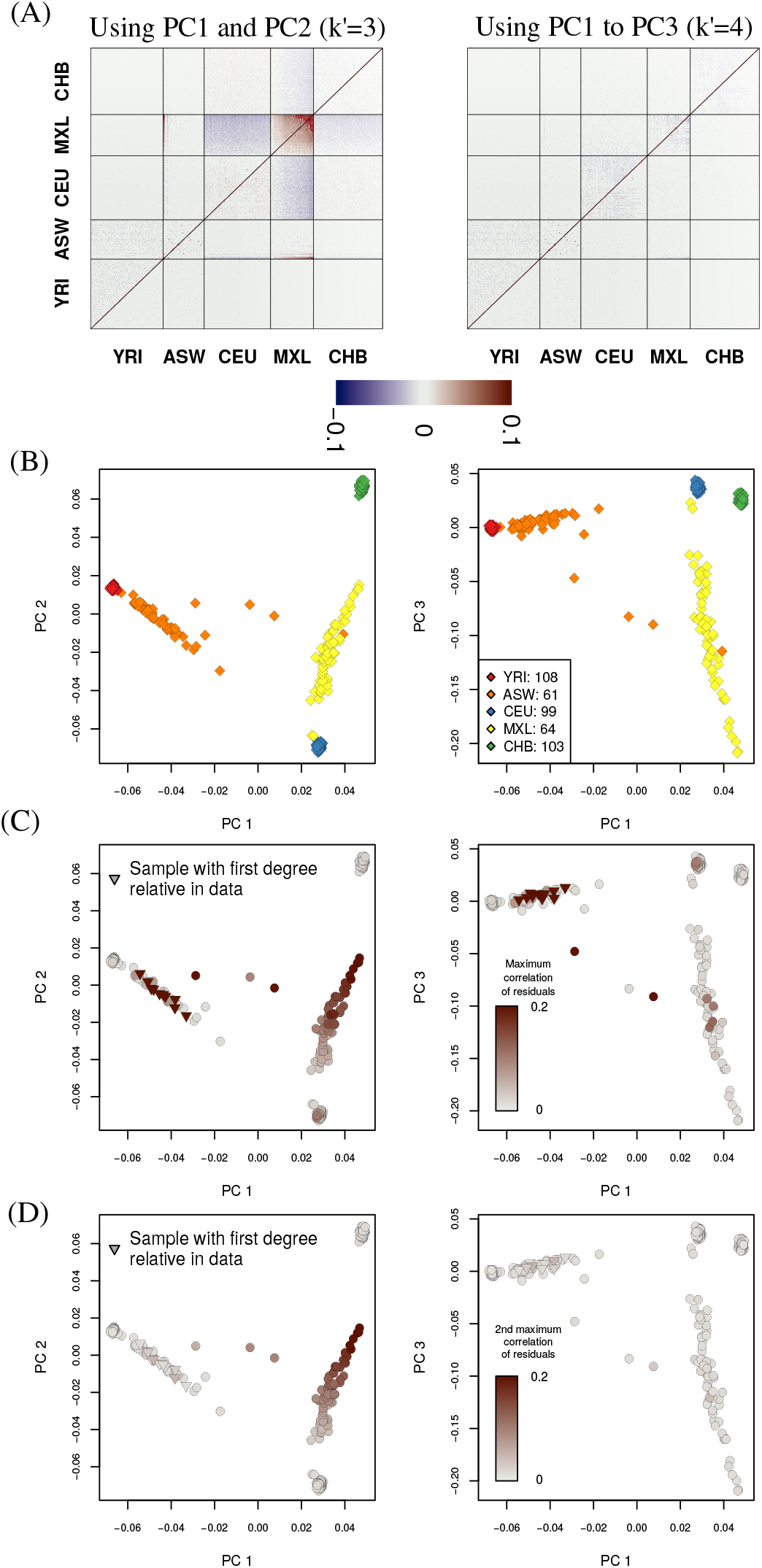
Population structure evaluated on human data from the 1000 Genomes project based on centered genotypes (PCA 2). (A) The upper triangle in the plots shows the empirical correlation coefficient 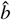 and the lower triangle shows the difference 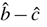. (B) Visualization of the three major principal components colored by the population of origin. (C) Visualization of the three major principal components colored by the maximum residual correlation coefficient an individual has with all remaining individuals. Individuals that have one or more first degree relative in the data are marked with a triangle. (D) Visualization of the three major principal components colored by the second maximum residual correlation coefficient an individual has with all remaining individuals. Individuals that have one or more first degree relative in the data are marked with a triangle.

We also evaluated the approach on the very diverse Human Genome Diversity Panel (HGDP) dataset (Cann *et al*. 2002). We used the compilation of the data described in (Patterson *et al*. 2012), and excluded non *Homo sapiens* samples, non-autosomal SNPs and SNPs with MAF < 0.05 in the selected samples. This resulted in keeping keeping in total 429,928 for 934 individuals from 53 populations. The results are shown in Figure S6. The correlation coefficients clearly show that on a continental scale the first two PCs places the Americas, east Asia and Oceania too close to each other. With 5 PCs, most of the differences between continents have been resolved. However, most populations still have a distorted placement in PC space compared to other populations within the same continental group. This is especially true within Africa, Oceania and the Americas, where the populations are genetically more distant to each other than what is reflected by the top 5 PCs. Most of negative correlation between populations is resolved with 17 PCs. However, not even 24 PCs are sufficient to accurately portrait the population structure of the 53 populations.

## Discussion

We have developed a novel approach to assess model fit of PCA and the admixture model based on structure of the residual correlation matrix. We have shown that it performs well for simulated and real data, using a suit of different PCA methods, commonly used in the literature, and the ADMIXTURE software to estimate model parameters. By assessing the residual correlation structure visually, one is able to detect model misfit and violation of modelling assumptions.

The model fit is assessed by comparing visually two matrices of residual correlation coefficients. The theoretical and practical advantage of our approach lie in three aspects. First, our approach is computationally simple and fast. Calculation of the two residual correlation matrices and their difference is computationally inexpensive. Secondly, our approach provides a unified approach to model fitting based on PCA and clustering methods (like ADMIXTURE). In particular, it provides simple means to assess the adequacy of the chosen number of top principal components to describe the structure of the data. Assessing the adequacy by plotting the principal components against each other might lead to false confidence. In contrast, our approach exposes model misfit by plotting the difference between two estimated matrices of the residual correlation coefficients. Thirdly, it comes with theoretical guarantees in some cases. These guarantees are further back up by simulations in cases, we cannot provide theoretical validity. Finally, our approach might be adapted to work on NGS data without estimating genotypes first, but working directly on genotype likelihoods.

## Data availability

The data sets used in this study are all publicly available, including simulated and real data. Information about the R code used to analyze and simulate data is available at https://github.com/popgenDK/evalPopStructure/. The variant calls for the 1000 Genomes Project data used are publicly available at ftp://ftp.1000genomes.ebi.ac.uk/vol1/ftp/release/20130502/ and the ones for the HGDP dataset are available at https://reich.hms.harvard.edu/sites/reich.hms.harvard.edu/files/inline-files/Data.tar. An R implementation of the method together with interactive tutorials are publicly available in https://github.com/popgenDK/evalPopStructure/.

## Acknowledgements

The authors are supported by the Independent Research Fund Denmark (grant number: 8021-00360B) and the University of Copenhagen through the Data+ initiative. SL acknowledges the financial support from the funding agency of China Scholarship Council. GGE and AA are supported by the Independent Research Fund Denmark (grant numbers: 8049-00098B and DFF-0135-00211B respectively).

## Appendix A

We first state the expectation and covariance matrix of *G*_*s⋆*_ and Π_*s⋆*_, respectively, under the given distributional assumptions,

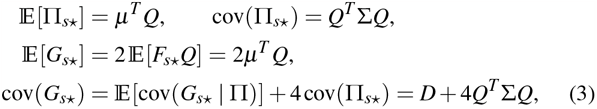

for *s* = 1, …, *m*, where

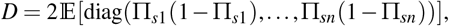

and

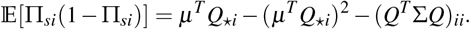

The unconditional columns *G*_*s⋆*_, *s* = 1, …, *m*, of *G* are independent random vectors by construction.

The above implies that

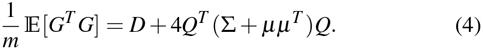

Auxiliary results are in appendix C.

### Lemma 4.

*The estimator* 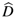 *is an unbiased estimator of D, that is*, 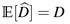. *Furthermore, it holds that* 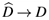 *as m* → ∞.

*Proof*. Conditional on Π_*si*_, using binomiality, we have 𝔼 [*G*_*si*_ (2− *G*_*si*_) |Π_*si*_] = 2Π_*si*_ (1 −Π_*si*_), and the first result follows. For convergence, note that *G*_*si*_ (2 −*G*_*si*_), *s* = 1, …, *m*, unconditionally, form a sequence of iid random variables with finite variance, hence the convergence statement follows from the strong Law of Large Numbers (Jacod and Protter 2004).

### Lemma 5.

*The estimator* 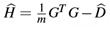 *is an unbiased estimator of H* = 4*Q* ^*T*^ (∑ + *μμ*^*T*^)*Q, that is*,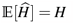. *Furthermore, it holds that* 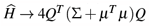 *as m* → ∞, *and*

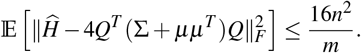

*Proof*. Unbiasedness follows from (4) and Lemma 4. Consider the (*i, j*)-th entry of 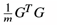, namely, 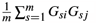. The sequence *G*_*si*_*G*_*s j*_, *s* = 1, …, *m*, is iid with finite variance, hence 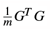 converges to 𝔼 [*G*^*T*^ *G*] as *m*→ ∞ by the strong Law of Large Numbers (Jacod and Protter 2004). Combined with Lemma 4 gives convergence of 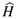 to *H* as *m*→ ∞.

It remains to prove the inequality. Define

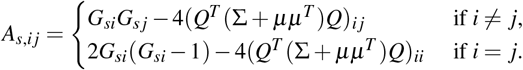

Then,

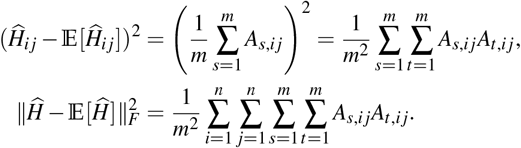

Using 𝔼 [*A*_*s,i j*_] = 0, independence of *A*_*s,i j*_ and *A*_*t,i j*_ for *s* ≠ *t*, and |*A*_*s,i j*_ | ≤ 4, we have

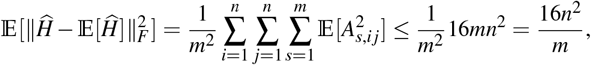

which proves the claim.□

The convergence result is also in Chen and Storey (2015, theorem 2). The second part provides the rate of convergence of 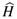 in the *L*^2^ - norm. Convergence is contingent on large *m*, rather than large *n*, and requires *m* to increase at least like the square of *n*.

**Proof of Theorem 1**. Since 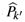 is assumed to be an orthogonal projection, that is, 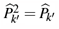 and 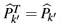, then also the limit is an orthogonal projection, 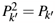 and 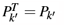.

Consider the empirical covariance 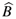. Define the variables *T*_*k*_′ = *G*(*I* − *P*_*k*_′) with 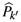 replaced by *P*_*k*_′, and the empirical covariance

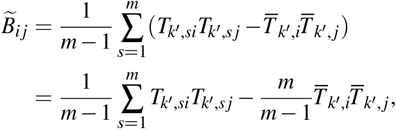

defined similarly to 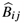, with 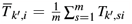. The sequences *T*_*k*_′_,*si*_*T*_*k*_′_,*s j*_, *s* = 1, 2, …, and *T*_*k*_′_,*si*_, *s* = 1, 2, …, are iid random variables, by the distributional assumptions on *G*. Furthermore, since *P*_*k*_′ is an orthogonal projection, then 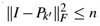 is bounded (Lemma 12). Therefore, also *T*_*k*_′_,*si*_ is bounded uniformly in *s, i* by 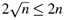.

Using boundedness, independence and the strong Law of Large Numbers (Jacod and Protter 2004),

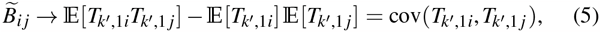

for *m* →∞, and cov(*T*_*k*_′_,1*i*_, *T*_*k*_′_,1 *j*_) = (*I* −*P*_*k*_′)(*D* + 4*Q* The latter equality follows from (4).

Consider 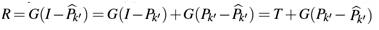. Hence,

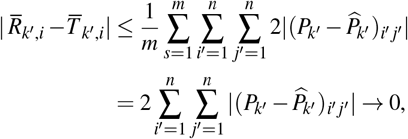

as *m* → ∞ by assumption of the theorem. It follows that 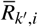 converges to 𝔼 [*T*_*k*_′_,1*i*_] as *m* → ∞. Furthermore,

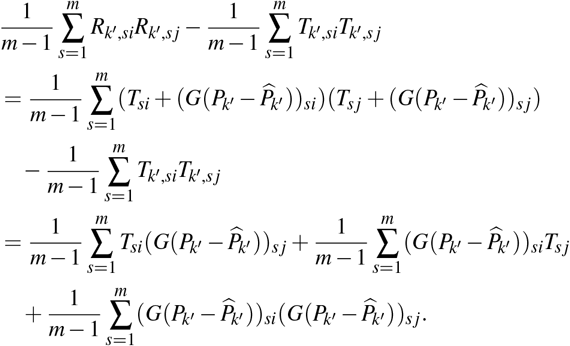

The absolute value of the first term in the last line above is bounded by

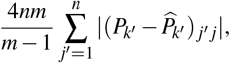

and similarly for the second term. The third is bounded by

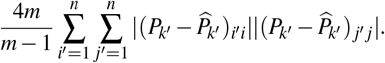

All three terms converge to zero as *m* → ∞, hence we conclude from (5) that 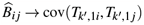 as *m* → ∞.

The result for the estimated covariance 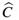 follows from convergence of 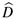 and by assumption of the theorem. The remaining part follows from the convergence of 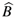 and 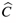. Note that *QP* = *Q*, hence the second equation holds. The last statement of the theorem follows directly.

**Proof of Theorem 2**. Consider *T*_*k*_ = *G*(*I* −*P*_*k*_) = *G*(1 −*P*), where *P* = *Q*^*T*^ (*QQ*^*T*^)^−1^*Q* is the projection onto the row space of *Q*. Then, *T*_*k*_ contains the residuals under multiple regression of the *m* rows of *G* on the *k* rows of *Q* (Box *et al*. 2005). Since *e* is in the row space of *Q*, then the sum of the residuals is zero for each *s* = 1, …, *m*: 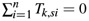 (the assumption that *e* is in the row space is equivalent to having an intercept in the regression model) (Box *et al*. 2005). We have, for *s* = 1, …, *m*,

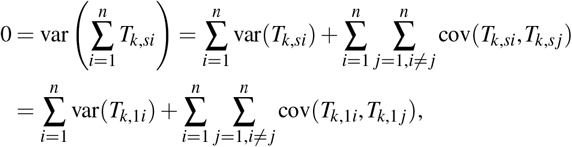

since the distribution of *T*_*k,si*_ is independent of *s*. From the proof of Theorem 6, it follows that 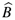 converges to cov(*T*_*k*,1*⋆*_) as *m* → ∞. Hence,

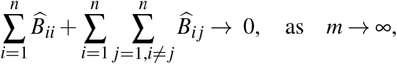

and the desired result follows by rearrangement.

If *Q* takes the given form, then the residuals under multiple regression are independent between compartments, as the projection is

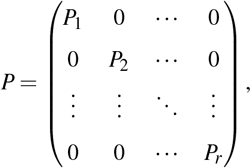

where 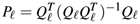 has dimension *n*_*ℓ*_ ×*n*_*ℓ*_. It follows that the computation above holds for each compartment. Finally, if *Q*_*ℓ*_ = (1 … 1), then the distribution of the random vector *T*_*k*,1*⋆*_ is exchangeable, resulting in

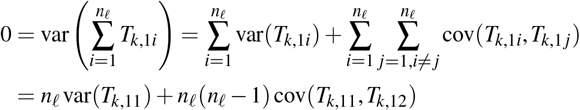

assuming the individuals in the *ℓ*-th compartment are numbered 1 to *n*_*ℓ*_. Rearranging terms and substituting 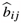 for the moments of *T*_*k,i⋆*_ yields the desired result.

**Proof of Theorem 3**. Consider *T*_*k*_ = *G*(*I*− *P*_*k*_) = *G*(1− *P*), where *P* = *Q*^*T*^ (*QQ*^*T*^)^−1^*Q* is the projection onto the row space of *Q*. If *Q*^1^ = (1 … 1), then the distribution of the random variables 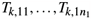 are exchangeable, resulting in

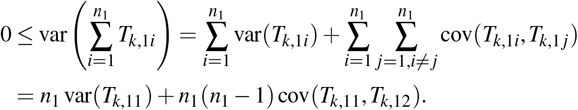

Rearranging terms and substituting 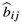 for the moments of *T*^*k,i⋆*^ yields the desired result.

**Proof of Theorem 6**. The convergence statement of the theorem is a special case of Theorem 9 in Appendix B. Take 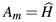 (that depends on the number of SNPs *m*, and the particular realization), *A* = *H*, and *k* = *k*^′^ in the theorem (*k* is used as a generic index in Theorem 9). Then, 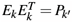 and 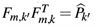, and the conclusion of Theorem 6 holds.

Convergence in Frobenius norm is equivalent to pointwise convergence (as *n* is fixed) 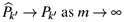 by definition.

If ∑ + *μμ*^*T*^ is positive definite, then it has rank *k*. As rank(*Q*) = *k* by assumption, it follows from Lemma 13 that rank(*H*) = *k*. Consequently, there are *k* positive eigenvalues of *H* and *λ*_*k*+1_ = 0, and the eigenvalue condition holds. Conversely, assume the eigenvalue condition holds. By definition rank(*H*) ≤ *k*. As *λ*_*k*_ *> λ*_*k*+1_ ≥0 by assumption, then also rank(*H*) ≥*k* and we conclude rank(*H*) = *k*. It follows that the rank of ∑ + *μμ*^*T*^ is *k*; consequently, it is positive definite.

If *k*′ = *k* = *n*, then 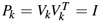 and *P* = *I* (as *k* = *n*), and *P*_*k*_ = *P*. So assume *k*′ = *k < n*. Since the eigenvalue condition is fulfilled, then from the above, we have rank(*Q*^*T*^ (∑ + *μμ*^*T*^)) = *k*, and Lemma 13 yields that the row space of *H* and *Q* agree. Similarly, we have 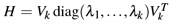 and Lemma 13 yields that the row space of *H* and 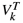 agree. This implies the row space of *Q* and 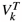 agree. Consequently, *P*_*k*_ = *Q*^*T*^ (*QQ*^*T*^)^−1^*Q* = *P*, and the statement holds.

**Proof of Theorem 7**. It follows trivially that *e* is an eigenvector of *H*_1_ with eigenvalue 0. If *D* has all entries positive, then it is positive definite and *D* + 4*Q*^*T*^ (∑ + *μμ*^*T*^)*Q* is also positive definite, hence has rank *n*. It follows from Lemma 13 that *H*_1_ has rank *n* − 1, hence *λ*_*n*−1_ *>* 0.

Similarly to the proof of Lemma 5 in Appendix B, one can show 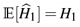 and 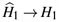 as *m* →∞, where *H*_1_ denotes the right hand side of (7). The remaining part of the theorem is proven similarly to Theorem 6.

**Proof of Theorem 8**. Note that *e* is an eigenvector of *H*_1_ with eigen-value 0. Consider an eigenvector *v* of *H*_1_, orthogonal to *e* with eigen-value *λ*. Then, the following two equations are equivalent,

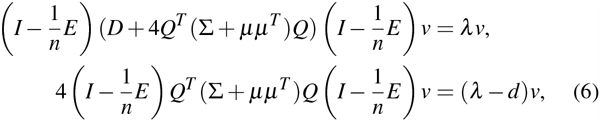

where it is used that *D* = *dI* and *v* ⊥ *e*. It shows that *v* is an eigenvector of 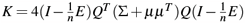 with eigenvalue *μ* = *λ* −*d*. Since *Q* has rank *k* and the vector *e* is in the space spanned by the rows of *Q*, then 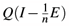 has rank *k* − 1. It follows that there are at most *k* −1 positive eigenvalues of *K*, that is, at most *k*− 1 eigenvalues of *H*_1_ such that *λ > d*. Furthermore, there are precisely *k* − 1 positive eigenvalues, provided ∑ + *μμ*^*T*^ is positive definite (Lemma 13). The remaining eigenvalues of *K* are zero, that is, the corresponding eigenvalues of *H*_1_ are *λ* = *d*.

Assume ∑ + *μμ*^*T*^ is positive definite, then by the above argument there precisely are *k* −1 eigenvalues of *H*_1_ such that *λ > d* with corresponding orthogonal eigenvectors *v*_1_, …, *v*_*k*− 1_. It follows from (6) that *v*_1_, …, *v*_*k* −1_ are in the space spanned by the rows of 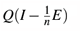, hence the eigenvectors are in the space spanned by the rows of *Q*. By assumption *e* is also in that row span. Hence, *v*_1_, …, *v*_*k*−1_, *e* forms an orthogonal basis of the row span of *Q*, as *Q* has rank *k*. Thus, *P*_*k*_ = *P*.

## Appendix B

In this appendix we give mathematical guarantees for correctness of the two approaches PCA 1 and PCA 2.

### Using PCA 1

In this particular case, convergence of 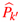 can be made precise. Define the matrix *H* = 4*Q*^*T*^ (∑ + *μμ*^*T*^)*Q*. Then, *H* is symmetric and positive semi-definite because ∑ and *μμ*^*T*^ both are positive semi-definite. Hence, *H* has non-negative eigenvalues. Furthermore, according to Lemma 5 in Appendix A, *H* converges to 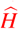 as *m* → ∞.

#### Theorem 6.

*Assume k*′≤ *k. Let λ*_1_ ≥… ≥*λ*_*n*_ ≥0 *be the eigenvalues of H, with corresponding orthogonal eigenvectors v*_1_, …, *v*_*n*_. *In particular, λ*_*k*+1_ = … = *λ*_*n*_ = 0, *as Q has rank k. Let P*_*k*_′ *be the orthogonal projection onto the span of v*_1_, …, *v*_*k*_′, *that is*,

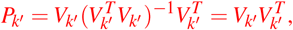

*where V*_*k*_′ = (*v*_1_, …, *v*_*k*_′).

*Assume k*′ = *n or λ*_*k*_′ *> λ*_*k*_′ _+1_, *referred to as the eigenvalue condition. Then*, 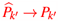 *as m* →∞. *If the eigenvalue condition is fulfilled for k*′ = *k, then P*_*k*_ = *P, that is, P*_*k*_ *is the orthogonal projection onto the span of the row vectors of Q. In particular, the eigenvalue condition is fulfilled for k*′ = *k if and only if* ∑ + *μμ*^*T*^ *is positive definite. The latter is the case if* ∑ *is positive definite*.

### Using PCA 2

The squared singular values in the SVD decomposition of *G*_1_ are the same as the eigenvalues of

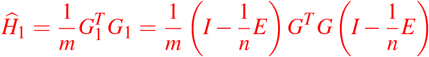

(Jolliffe 2002). We have

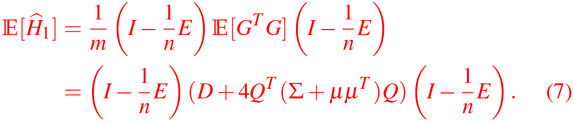

Let *H*_1_ denote the right hand side of (7).

#### Theorem 7.

*Let λ*_1_ ≥… ≥ *λ*_*n*_ *be the eigenvalues of H*_1_, *with corresponding orthogonal eigenvectors v*_1_, …, *v*_*n*_. *In particular, v*_*n*_ = *e and λ*_*n*_ = 0. *If D has all diagonal entries positive, then λ*_*n*−1_ *>* 0.

*Let k*′ ≤ *n and let P*_*k*_′ *be the orthogonal projection onto the span of v*_1_, …, *v*_*k*_′ _−1_, *e, that is*,

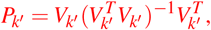

*where V*_*k*_′ = (*v*_1_, …, *v*_*k*_′ _−1_, *e*). *If k*′ = *n or λ*_*k*_′ *> λ*_*k*_′ _+1_, *then* 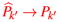 *as m* → ∞.

There are no guarantees that for *k*′ = *k*, we have *P*_*k*_ = *P* and that the difference between 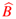 and 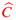 converges to zero for large *m*. However, this is the case under some extra conditions, and appears to be the case in many practical situations, see the ‘Results’.

#### Theorem 8.

*Assume D* = *dI for some d >* 0. *Furthermore, assume the vector e is in the row space of Q (this is, for example, the case if the admixture proportions sum to one for each individual). Then, λ*_*k*_ = … = *λ*_*n* −1_ = *d, and λ*_*n*_ = 0.

*If* ∑ + *μμ*^*T*^ *is positive definite, then λ*_*k*+1_ *> λ*_*k*_ *and P*_*k*_ = *P, where P*_*k*_ *is as in Theorem 6. As a consequence, with k*′ = *k in Theorem 1*, 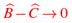 *as m* → ∞.

## Appendix C

### Theorem 9.

*Let A*_*m*_ *be a sequence of symmetric n×n-matrices that converges to a symmetric n* ×*n-matrix A in the Frobenius norm, that is* ∥*A*_*m*_ −*A*∥ _*F*_→ 0, *as m*→ ∞. *Let λ*_1_ ≥… ≥*λ*_*n*_ *be the eigenvalues of A (with multiplicity, and not necessarily non-negative). Let k*≤ *n be given and assume either k* = *n or λ*_*k*_ *> λ*_*k*+1_. *Furthermore, let e*_1_, …, *e*_*k*_ *be orthogonal eigenvectors corresponding to the eigenvalues λ*_1_, …, *λ*_*k*_, *respectively, and let f*_*m*,1_, …, *f*_*m,k*_ *be orthogonal eigenvectors corresponding to the k largest eigenvalues of A*_*m*_ *(with multiplicity). Then, the orthogonal projection onto the span of f*_*m*,1_, …, *f*_*m,k*_ *converges to the orthogonal projection onto the span of e*_1_, …, *e*_*k*_ *in the Frobenius norm. That is, define E*_*k*_ = (*e*_1_, …, *e*_*k*_) *and F*_*m,k*_ = (*f*_*m*,1_, …, *f*_*m,k*_), *then* 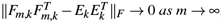.

*Proof*. If *k* = *n*, then 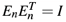 and 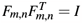, and the statement is trivial. Hence, assume *k < n*. Let *e*_1_, …, *e*_*n*_ be eigenvectors of *A* corresponding to eigenvalues *λ*_1_, …, *λ*_*n*_, respectively. Let *f*_*m*,1_, …, *f*_*m,n*_ be the eigenvectors of *A*_*m*_ corresponding to the eigenvalues *μ*_*m*,1_≥ … ≥*μ*_*m,n*_. All eigenvectors can be asssumed to be orthonormal.

As 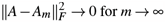, then every entry of *A*_*m*_ converges to the corresponding entry of *A*. Consequently, the characteristic function of *A*_*m*_ converges to that of *A*, and the eigenvalues of *A*_*m*_ converges to those of *A*, that is, *μ*_*m, j*_→ *λ*_*j*_ for *j* = 1, …, *n*, and *m* →∞. Let *T*_*m*_ be such that *E*_*n*_ = *F*_*m,n*_*T*_*m*_. As *E*_*n*_ and *F*_*m,n*_ are orthogonal matrices, hence also *T*_*m*_ is orthogonal. Applying Lemma 10 in the first and third line gives

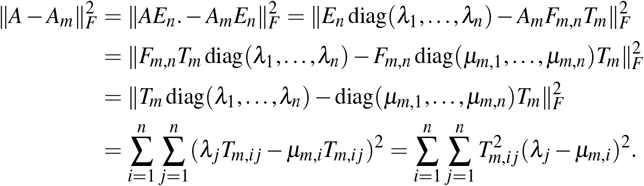

By assumption, *λ*_*k*_ > *λ*_*k*+1_. Hence, by convergence of eigenvalues, for *j* ≤ *k, i* ≥ *k* + 1, or *j* ≥ *k* + 1, *i* ≤ *k*, we have *T*_*m,i j*_ → 0 as *m* → ∞.

Furthermore,

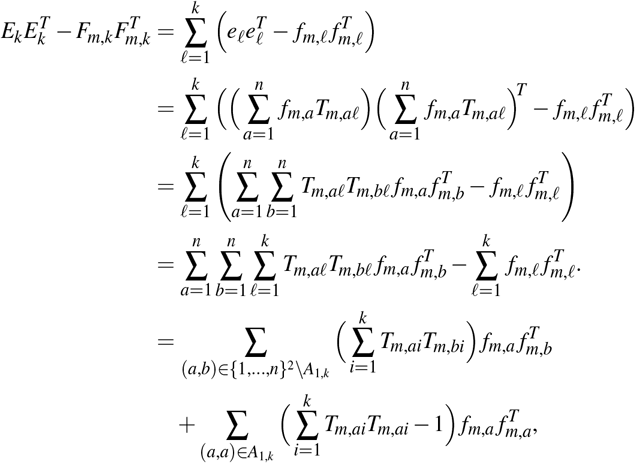

where *A*_*i, j*_ = {(*a, a*) : *i* ≤ *a* ≤ *j*}.

From Lemma 11, we have 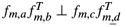 for (*a, b*) ≠ (*c, d*) in the Frobenius inner product. Moreover, 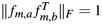 for all *a, b*. Hence,

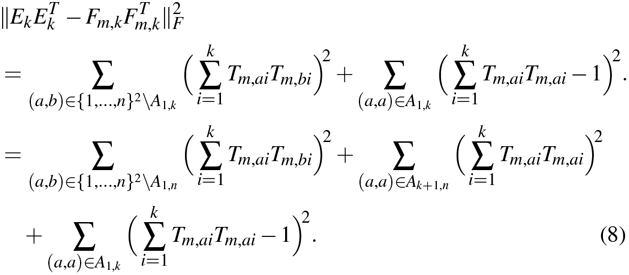

As noted above, *T*_*m,i j*_ → 0 as *m* → ∞ for *j* ≤ *k, i* ≥ *k* + 1, or *j* ≥ *k* + 1, *i* ≤ *k*. Using this and orthogonality of *T*_*m*_ gives

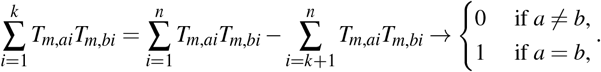

Inserting into (8) results in 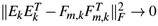, as *m* → ∞.□

### Lemma 10.

*Let A be an a* × *b matrix. Let U be a b* × *b orthogonal matrix and V an a* × *a orthogonal matrix. Then*,

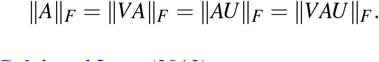

*Proof*. See Golub and Loan (2013).□

### Lemma 11.

*Let w, x, y, z* ∈ ℝ^*b*^. *For a* × *b-matrices A and B, let* 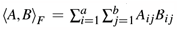 *be the Frobenius inner product of A*\*and B, an*l*d let* ⟨ ·, · ⟩ *be the standard inner product on ℝ*^*b*^. *Then*,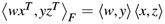. *In particular*, 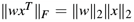 *and wx*^*T*^ ⊥ *yz*^*T*^ *if w* ⊥ *y or x* ⊥ *z. Proof*. Note that

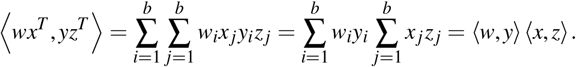

Hence, if leither *w* ⊥ *y* or *x* ⊥ *z*, then *wx*^*T*^ ⊥ *yz*^*T*^, and 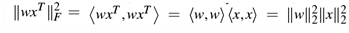, such that ∥*wx*^*T*^ ∥ _*F*_ = ∥*w*∥_2_∥*x*∥_2_.

### Lemma 12.

*Let v*_1_, …, *v*_*ℓ*_ *be linearly independent vectors. An orthogonal projection matrix on span*(*v*_1_, …, *v*_*ℓ*_) *has Frobenius norm* 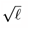.

*Proof*. We may assume that *v*_1_, …, *v*_*ℓ*_ are orthonormal. Then, we can write the projection matrix as 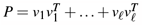. By Lemma 11, 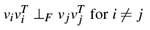. So, again by Lemma 11, 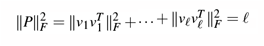.

### Lemma 13.

*Let A be an a*× *b matrix and B an b* ×*c matrix, both of rank b, such that a, c* ≥*b. Let C* = *AB. Then, C is of rank b, and the row space of C coincides with the row space of B*.

*Proof*. First we show that rank(*C*) = *b*. Note that *A* has *b* linearly independent rows 1 ≤ *i*_1_ *<* … *< i*_*b*_ ≤ *b*, and *B* has *b* linearly independent columns 1 ≤ *j*_1_ *<* … *< j*_*b*_ ≤ *b*. Let *Ã* and 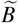 be the *b* × *b* matrices with 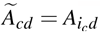 and 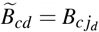. Then *Ã* and 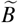 are invertible matrices. Hence, also 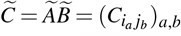 is invertible and has rank *k*. It follows that *C* has rank *k*. As *Ã* is invertible, then the span of the rows of *AB* is equal to the span of the rows of *B*. That is, the span of the rows of *AB* is equal to the span of the rows of *B*.□

